# Retrieval of the Complete Coding Sequence of the UK-Endemic Tatenale Orthohantavirus Reveals Extensive Strain Variation and Supports its Classification as a Novel Species

**DOI:** 10.1101/844340

**Authors:** Joseph G. Chappell, Theocharis Tsoleridis, Okechukwu Onianwa, Gabby Drake, Ian Ashpole, Phillipa Dobbs, William Edema, Frederick Kumi-Ansah, Malcolm Bennett, Rachael E Tarlinton, Jonathan K Ball, C. Patrick McClure

## Abstract

Hantaviruses are a diverse group of single-stranded, negative-sensed RNA viruses, known to cause sporadic outbreaks of potentially fatal human disease. To date, only two Orthohantavirus species have been detected in the UK - Seoul virus and Tatenale. Whilst Seoul is known to be pathogenic in humans, only partial fragments of Tatenale have been recovered, precluding any accurate analysis of its phylogeny or potential pathogenicity. To overcome this shortfall we used a degenerate primer PCR method to identify Tatenale-infection in rodents living in two separate locations in the UK. PCR positive samples were then subjected to either unbiased high-throughput sequencing or overlapping PCR product sequencing to recover the complete coding sequence of the Tatenale virus. This analysis provided in-depth insight into the evolutionary origins of this recently identified UK *Orthohantavirus* and unequivocally showed that Tatenale virus meets the established criteria for classification as a novel species. Crucially, our data will facilitate *in vitro* investigation into the zoonotic potential of Tatenale virus.

## Main Text

Orthohantaviruses are a diverse genus of RNA viruses, belonging to the *Bunyavirales* order. They primarily circulate in rodents, in which infection is persistent and asymptomatic. Several species are capable of transmission into humans through the inhalation of aerosolised excreta. Infection can lead to hantavirus fever (HF) [1], with severity ranging from sub-clinical to fatal. Four species are known to cause HF in Europe: Seoul (SEOV), Dobrava-Belgrade (DOBV), Tula (TULV) & Puumala (PUUV) [2]; only one of which, SEOV, has been detected in the UK [3].

HF has been reported sporadically in the United Kingdom [4], although the causative species could not be identified due to cross-reactivity of the serological assays used to diagnose the Orthohantavirus infections. In addition, SEOV RNA was detected in 2011 & 2012 in brown rats (*Rattus norvegicus*) that were epidemiologically linked to 2 HF cases [5]. Furthermore, a novel vole-associated Hantavirus related to TULV & PUUV, Tatenale virus (TATV), was identified in North West England in 2013 [6] and again in Northern England in 2017 [7]. However, fragments of less than 400 nucleotides were retrieved for only two of the three genomic segments, meaning that phylogenetic analysis of this virus was limited. In 2019, an orthohantavirus was detected in German field voles, Traemersee virus (TRAV), and was thought to be a strain of Tatenale virus. However, the aforementioned paucity of published TATV sequence data has precluded any accurate comparison between TATV and TRAV [8].

To better understand Tatenale virus prevalence and phylogeny, we performed in-depth sampling and analysis of various rodents living in the UK. Rodents were caught at two sites in the United Kingdom: Leicestershire (Site 1) and Cheshire (Site 2). Seventy-two rats (*R. norvegicus*), 224 mice (*Mus musculus*) and 12 field voles (Microtus agrestis) were collected from Site 1 between May 2013 and October 2014 [9]. Eight rats, 119 field voles, 93 wood mice (*Apodemus flavicollis*) and 3 bank voles (*Myodes glareolus*) from site 2 between June 2013 and July 2016.

Lung and kidney tissues were collected and RNA extracted using GenElute™ Mammalian Total RNA Miniprep Kit (Sigma Aldrich). Two-step RT-PCR was performed on the samples, using a degenerate primer pair, designed to target a 178bp region of the L segment of all known Hantaviruses (Supplement). One field vole from site 1 (8.3%) and 12 field voles from site 2 (10%) were Hantavirus PCR positive and were rescreened using a second degenerate primer pair [10] targeting a larger fragment of L. Amplicons were subsequently sent for Sanger sequencing (SourceBioscience, Nottingham). BLAST homology searches of the resulting sequence data showed that both were related to Tatenale virus; we named the strain from site 1 ‘Norton-Juxta’ and the strain from site 2 ‘Upton-Heath’, reflecting the locations at which they were found.

To retrieve the complete genome, Norton-Juxta was subjected to unbiased high-throughput sequencing (uHTS). NEBNext® rRNA Depletion Kit (New England Biolabs) was used to remove host ribosomal RNA and then prepared for uHTS using the NEBNext® Ultra™ II RNA Library Prep Kit (New England Biolabs). The sample was then sequenced using an Illumina HiSeq 4000 (Source Bioscience, Nottingham), generating 27 Million paired reads. Paired reads were merged and trimmed using Geneious R11 and mapped to the reference sequences for PUUV small (S), medium (M) & L segments. The completed CDS of Norton-Juxta was used as a reference to design a series of PCR primer sets to amplify the CDS of the Upton-Heath strain.

Full-length coding sequence (CDS) and partial untranslated region (UTR) was retrieved for each of the three segments of the Norton-Juxta strain; the CDS was 1303bp for S (GenBank accession number MK883757), 3447bp for M (MK883759) and 6465bp for L (MK883761). Using the PCR primer walking approach, complete CDS was retrieved for S (MK883756) and L (MK883760) and near-complete CDS of the M (MK883758) for the Upton-Heath strain. The two strains were 94.1%, 91.3% and 90.6% similar at the nucleotide level, across the S, M and L segments, respectively. The original phylogenies of B41 and Kielder strains of TATV were based on <400bp fragments of the L and S or L segment, respectively [6, 7]. Analysis of the corresponding L and S sequences (Supplementary figure 1A-B) showed that both the Norton-Juxta and Upton-Heath strains clustered with TATV. Within the partial S segment phylogeny, the Upton-Heath and Norton-Juxta strains were 99% and 94% similar to B41. For the partial L segment, the Upton-Heath and Norton-Juxta strains were 87% and 94% similar to the B41, respectively, and 86% similar to the Kielder strain.

Further analysis of the L (Figure 1, A), M (Figure 1, B) and S (Figure 1, C) segments of the novel UK strains, together with representative sequences of globally sampled vole Hantaviruses shows that each of the segments cluster with TRAV with 100% bootstrap support. TRAV was the closest related orthohantavirus, with the CDS of each of the Norton-Juxta and Upton-Heath segments showing a comparable similarity. TRAV was 82.7%/96.8% (Nt/AA) similar to Norton-Juxta and 83%/96.5% similar to Upton-Heath across the S segment, 79.8%/94.2% and 80.8%/94.3% to Norton-Juxta and Upton-Heath across the M segment and 81.5%/96.4% similar to both TATV strains across the L segment. Pairwise evolutionary distance (PED) analysis of the concatenated S and M segments of Norton-Juxta and other vole-borne Orthohantaviruses showed values of between 0.12 and 0.27; comparison of the PED values between Norton-Juxta and TRAV were 0.05.

**Figure 1.**
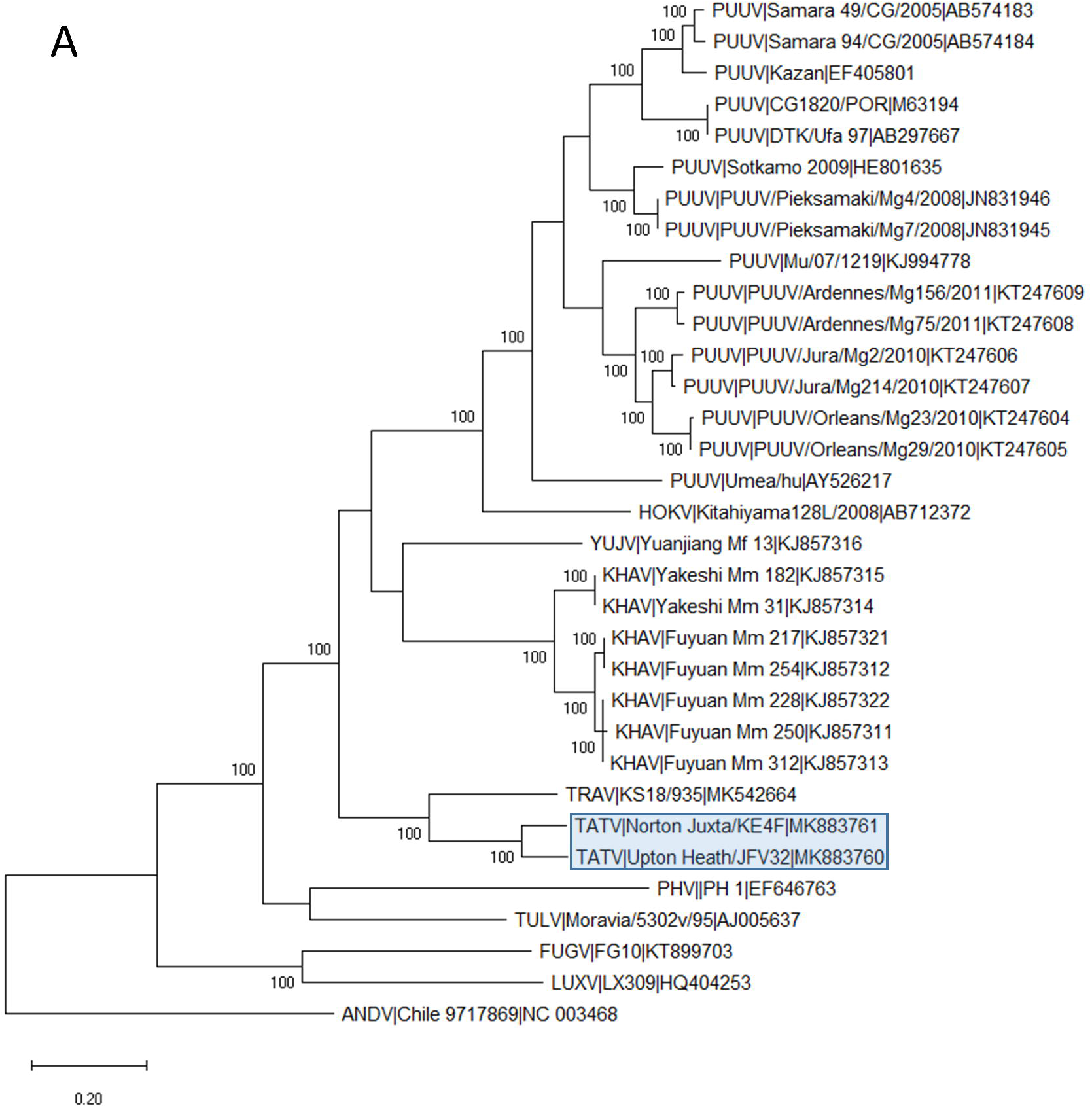
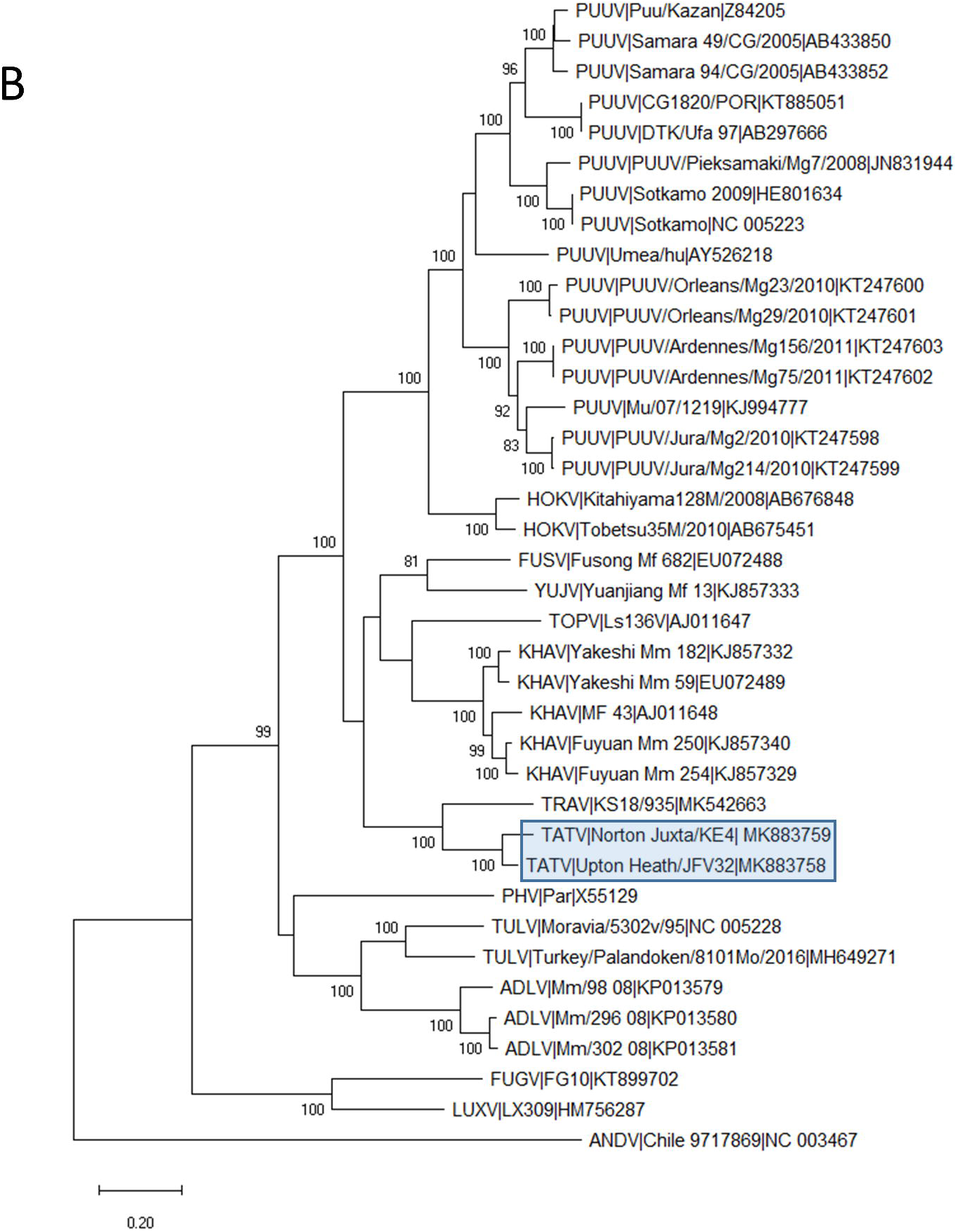
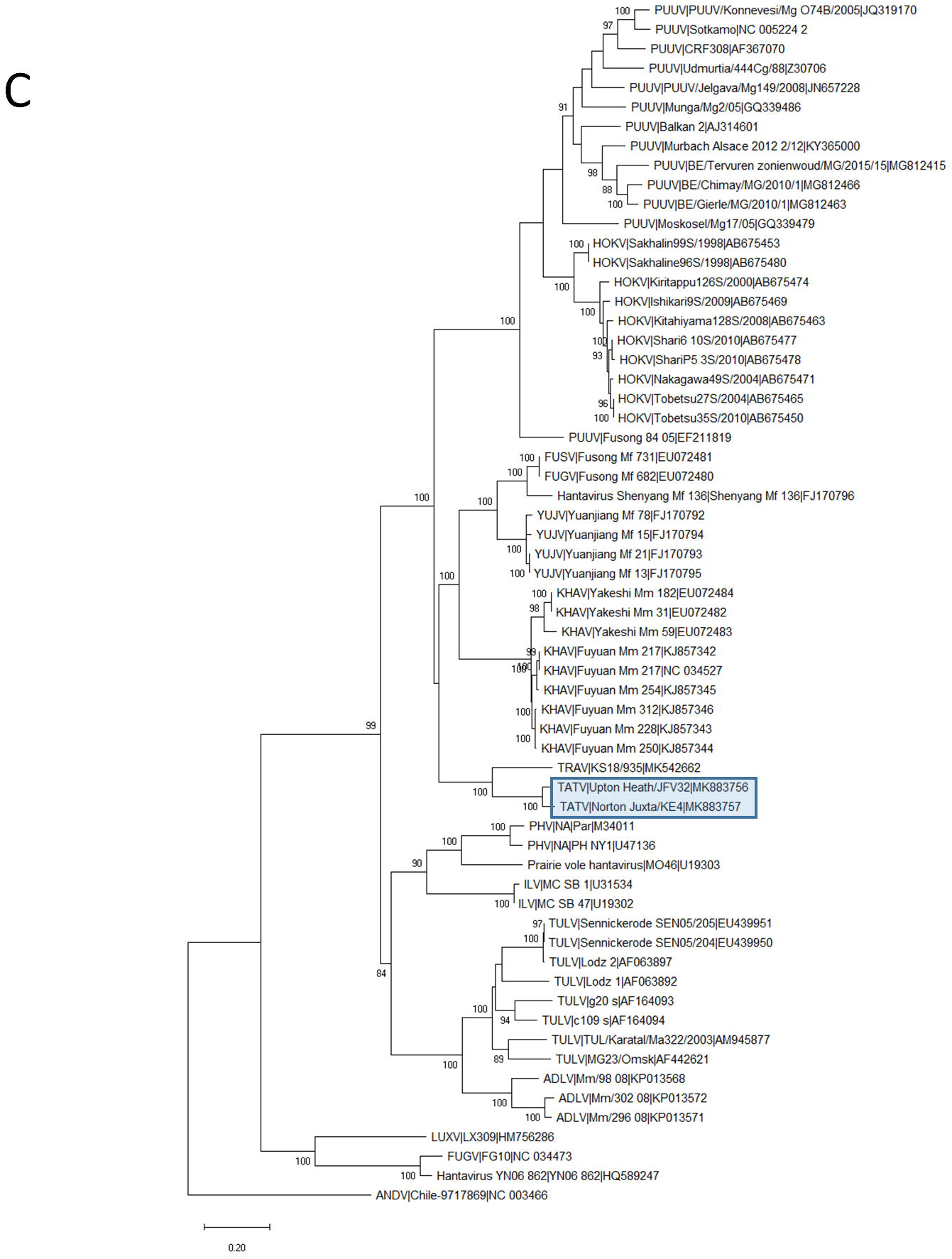
Phylogenetic relationship of Tatenale virus with other vole-associated orthohantavirus species. Representative complete coding sequences were retrieved for each segment; L (A), M (B) and S (C). Maximum Likelihood trees were created with a GTR+G+I model, using MEGAX software. Branch lengths are drawn to a scale of nucleotide substitutions per site. L and S trees were based on full-length sequences, while the M segment tree was based on the available sequence for the partial Upton-Heath strain. Numbers above individual branches show bootstrap support after 1000 replicates. Tatenale virus strains are highlighted with a blue box. Sequences are shown with the species name, strain name and the GenBank accession number. PUUV, Puumala virus; HOKV, Hokkaido virus; FUSV, Fusong virus; YUJV, Yuanjiang virus; KHAV, Khabarovsk virus; TOPV, Topografov virus; TATV, Tatenale virus; TRAV, Traemmersee virus; PHV, Prospect Hill virus; ILV, Isla Vista virus; TULV, Tula virus; ADLV, Adler virus; LUXV, Luxi virus; FUGV, Fugong virus; ANDV, Andes virus

This is the first reported recovery of full coding sequences for TATV. Species demarcation criteria of >7% AA divergence across S and M segments [11], as well as stricter criteria of a PED of lower than 0.1 in the concatenated S and M segments [12], have been suggested. As the PED values between the complete Norton-Juxta strain of TATV and TRAV is below the 0.1 speciation threshold, both viruses are members of the same viral species, as was hypothesised by Jeske *et al* [8].

These findings extend the known range of TATV into central England and further strengthens the evidence of *M. agrestis* as a primary reservoir. TATV, or a TATV-like virus, has been suggested as a possible causative agent of HF in the United Kingdom [7]. However, the paucity of sequence data has precluded significant investigation. Recovery of full-length CDS will allow for *in vitro* studies into the zoonotic potential of the virus

## References

1. Clement J, Maes P, Van Ranst M. Hemorrhagic Fever with Renal Syndrome in the New, and Hantavirus Pulmonary Syndrome in the old world: Paradi(se)gm lost or regained? Virus Research 2014;187:55–58.

2. Heyman P, Ceianu C, Christova I, Tordo N, Beersma M, et al. A five-year perspective on the situation of haemorrhagic fever with renal syndrome and status of the hantavirus reservoirs in Europe, 2005-2010. Eurosurveillance;16. http://www.eurosurveillance.org/ViewArticle.aspx?ArticleId=19961 (2011, accessed 10 March 2016).

3. Murphy EG, Williams NJ, Bennett M, Jennings D, Chantrey J, et al. Detection of Seoul virus in wild brown rats (Rattus norvegicus) from pig farms in Northern England. Veterinary Record 2019;184:525–525.

4. Watson AR, Irving WL, Ansell ID. Playing in a scrapyard and acute renal failure. Lancet 1997;349:1446.

5. Jameson LJ, Logue CH, Atkinson B, Baker N, Galbraith SE, et al. The continued emergence of hantaviruses: isolation of a Seoul virus implicated in human disease, United Kingdom, October 2012. Eurosurveillance 2013;18:20344.

6. Pounder KC, Begon M, Sironen T, Henttonen H, Watts PC, et al. Novel Hantavirus in field vole, United Kingdom. Emerging Infectious Diseases 2013;19:673–675.

7. Thomason AG, Begon M, Bradley JE, Paterson S, Jackson JA. Endemic Hantavirus in Field Voles, Northern England. Emerging infectious diseases 2017;23:1033–1035.

8. Jeske K, Hiltbrunner M, Drewes S, Ryll R, Wenk M, et al. Field vole-associated Traemmersee hantavirus from Germany represents a novel hantavirus species. Virus Genes 2019;1–6.

9. Tsoleridis T, Onianwa O, Horncastle E, Dayman E, Zhu M, et al. Discovery of Novel Alphacoronaviruses in European Rodents and Shrews. Viruses 2016;8:84.

10. Klempa B, Fichet-Calvet E, Lecompte E, Auste B, Aniskin V, et al. Hantavirus in African wood mouse, Guinea. Emerging infectious diseases 2006;12:838–40.

11. ICTV. ICTV Ninth Report; 2009 Taxonomy Release. https://talk.ictvonline.org/ictv-reports/ictv_9th_report/negative-sense-rna-viruses-2011/w/negrna_viruses/205/bunyaviridae (accessed 7 March 2019).

12. Maes P, Klempa B, Clement J, Matthijnssens J, Gajdusek DC, et al. A proposal for new criteria for the classification of hantaviruses, based on S and M segment protein sequences. Infection, Genetics and Evolution 2009;9:813–820.

