## Supplement for "Retrieval of the Complete Coding Sequence of the UK-Endemic Tatenale Orthohantavirus Reveals Extensive Strain Variation and Supports its Classification as a Novel Species"

**Pan-Hantavirus PCR Assay;**

HanSemiF: GAATATATATCNTAYGGDGGDGA

HanSemiR: CTGGTGACCAYTTNGTNGCAT

0.5μl of cDNA was added to reactions of 1.25μl Mg2+ free 10x PCR buffer supplemented with 0.06μl of HotStarTaq DNA Polymerase (Qiagen, Hilden, Germany), 0.5μl 10mM dNTP (Sigma Aldrich), 0.5μl of forward and reverse primer (10 Pmol/μl), and 9.19μl of water for a total volume of 12.5μl. Cycle conditions were 95°c for 15 minutes, 55 cycles of 94°c, 51°c and 72°c for 20 seconds each, followed by 72°c for 10 minutes.

**Phylogenetic Analysis**

Nucleotide sequence for each segment of TATV Norton Juxta & Upton Heath was aligned with full-length coding sequence for representative *Arvicolinae* associated Orthohantaviruses and a non-*arvicolinae* orthohantavirus (Andes Virus) outgroup, using the MUSCLE function in MEGAX [1]. MEGAX was then used to find the best-fit substitution model for each alignment of sequences, the model with the lowest Bayesian Information Criterion scores were considered the most appropriate.

**References**

[1]	Kumar S, Stecher G, Li M, et al. MEGA X: Molecular Evolutionary Genetics Analysis across Computing Platforms. Mol Biol Evol. 2018;35:1547–1549.

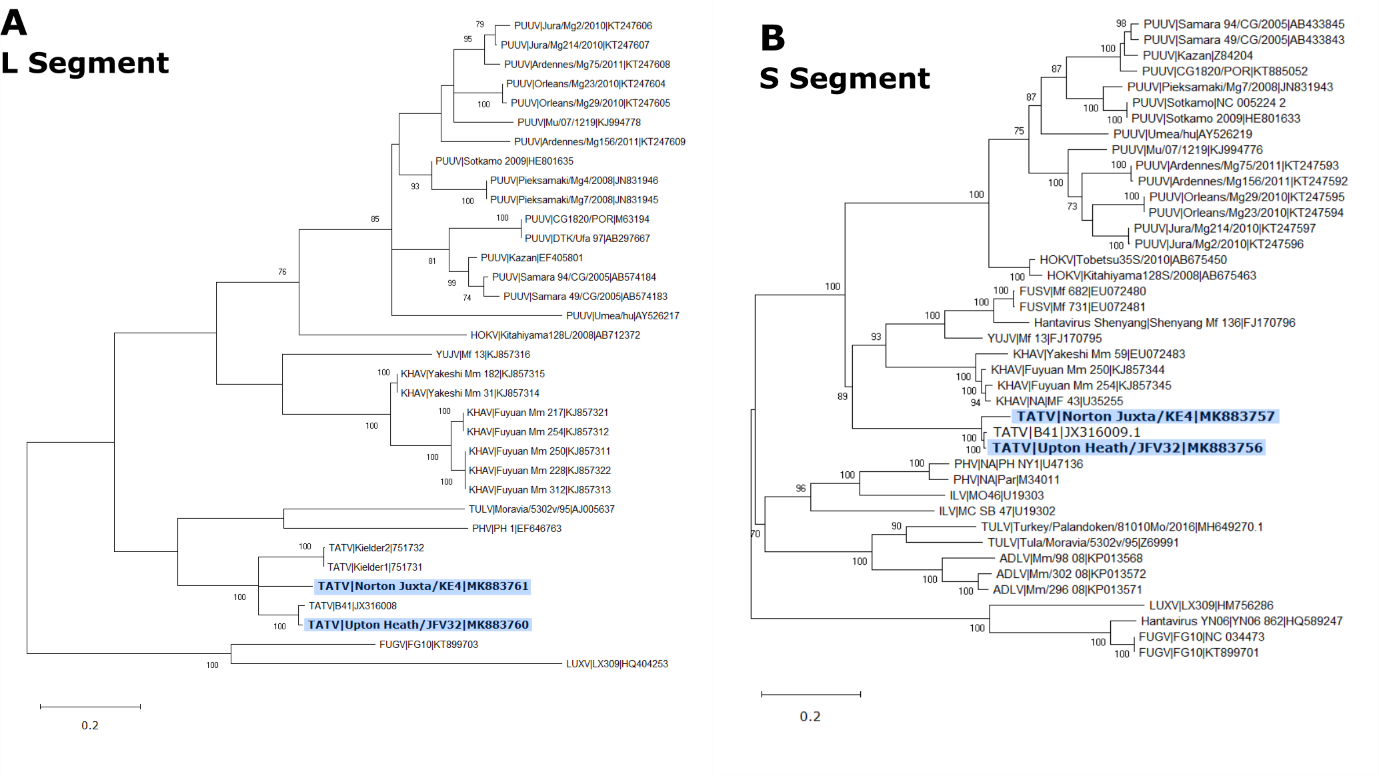
**Supplemental Figure**

Supplement Figure 1: Phylogenetic relationship of Tatenale virus with previous Tatenale isolates and other vole-associated orthohantavirus species. Representative partial sequences were obtained for L (A) and S (B) segments. Maximum Likelihood trees were created using the T92+G+I model, branch lengths are drawn to a scale of nucleotide substitutions per site. Numbers above individual branches show bootstrap support after 100 replicates. Tatenale virus strains are highlighted in boldface and a blue box. Sequences are shown with species name, strain name and the GenBank accession number. PUUV, Puumala virus; HOKV, Hokkaido virus; FUSV, Fusong virus; YUJV, Yuanjiang virus; KHAV, Khabarovsk virus; TATV, Tatenale virus; PHV, Prospect Hill virus; ILV, Isla Vista virus; TULV, Tula virus; ADLV, Adler virus; LUXV, Luxi virus; FUGV, Fugong virus.

**Table**

| Species | S | M | L |
| --- | --- | --- | --- |
|  | Norton-Juxta | | |
| Khabarovsk | 79.2 (89.4) | 76.4 (87.5) | 77.9 (90.9) |
| Yuanjiang | 79.2 (88.5) | 75.3 (86.5) | 77.7 (90.4) |
| Fusong | 78.7 (88.2) | 75.2 (85.9) | -* |
| Puumala | 77.9 (87.8) | 74.8 (84.7) | 77.9 (88.1) |
| Hokkaido | 78.3 (87.5) | 75.5 (84.4) | 76.8 (88.5) |
|  | Upton Heath | | |
| Khabarovsk | 79.9 (88.9) | 77.1 (87.8) | 78 (90.7) |
| Yuanjiang | 78.9 (88.2) | 75.7 (86.5) | 77.7 (89.6) |
| Fusong | 78.9 (88) | 76 (86.2) | -* |
| Puumala | 78.4 (87.8) | 75.5 (84.6) | 77.6 (87.5) |
| Hokkaido | 79 (87.8) | 75.7 (84.4) | 76.7 (87.9) |

Table 1: Similarity of Norton Juxta and Upton Heath strains of Tatenale virus to the closest related strain of the most related species at nucleotide (Amino Acid) level. *Indicates no complete sequence data available
